# What a stimulus predicts, not what it depicts, determines striatal reward signals

**DOI:** 10.64898/2026.05.10.724107

**Authors:** Nicola Sambuco, Francesco Versace

## Abstract

Free viewing of emotional pictures activates motivational circuits as a function of arousal and valence, yet, because no real outcomes are delivered, anticipatory wanting and consummatory liking cannot be dissociated. Gustatory neuroimaging paradigms deliver actual rewards but use impoverished, affectively neutral cues that do not engage natural selective attention. We bridged these paradigms by presenting emotional images alongside food photographs that either predicted juice delivery (Food+) or did not (Food−), all within a single fMRI session. On each Food+ trial, participants indicated in real time whether they wanted the juice, enabling a within-subject dissociation of anticipatory from consummatory signals. Nucleus accumbens showed a large and selective response to Food+ cues that exceeded activation to both pleasant (erotica) and unpleasant (mutilation) high-arousal images, establishing that mesolimbic engagement tracked outcome prediction rather than emotional arousal, affective valence, or visual content. A temporal dissociation further revealed that nucleus accumbens carried the dominant anticipatory signal during the cue period, while ventromedial prefrontal cortex carried the dominant outcome-period signal at juice delivery, a pattern consistent with the wanting and liking distinction. Representational similarity analysis confirmed that outcome prediction, rather than emotional arousal, affective valence, or visual category, was the dominant organizing principle of the multivariate neural response across the full region-of-interest network. Together, these findings show that whether a visual stimulus engages reward circuitry depends less on what it depicts than on what it predicts, and provide a framework for studying individual differences in appetitive motivation and cue-induced eating.

**Significance statement:** Whether a food image engages brain reward circuitry depends not on what it depicts, but on what it predicts. We scanned participants while they viewed food images that either preceded a real opportunity to receive juice or did not, alongside erotic, threatening, and neutral scenes. Nucleus accumbens, a core reward region, responded selectively to food images predicting juice, with a response that exceeded even the response to erotic images. Ventromedial-prefrontal cortex, by contrast, tracked actual juice receipt, dissociating anticipation from consumption. Across a ten-region network, learned reward prediction, rather than emotional arousal, valence, or visual category, organized the neural response. These findings establish a human neuroimaging paradigm for studying how cue-driven motivation goes awry in obesity, addiction, and compulsive eating.

## Introduction

Stimuli that signal potential threat or reward engage a broad neural network that prioritizes their processing over neutral input (Lang & Davis, 2006). This principle, natural selective attention (Bradley, 2009), has been demonstrated extensively using emotionally relevant pictures: images depicting erotica, threat, or mutilation reliably activate visual cortex, amygdala, medial prefrontal cortex, and inferior frontal gyrus, with activation scaling with motivational significance rather than valence alone (Sabatinelli et al., 2011). These effects replicate even in small samples with high-arousal stimuli (Sabatinelli et al., 2023; Sambuco, 2022), and are disrupted transdiagnostically in psychiatric conditions involving affective dysregulation (McTeague et al., 2020; Sabatinelli et al., 2005; Sambuco, Bradley, Herring, Hillbrandt, et al., 2020).

A critical limitation of this paradigm, however, is that the free viewing picture paradigm cannot distinguish incentive salience, the motivational wanting attributed to reward-predictive cues through learning about their relationship with a significant outcome, which biases attention and drives goal-directed approach behavior (Berridge et al., 2009; Berridge & Robinson, 2016). Throughout, wanting and liking refer to incentive salience and hedonic impact, respectively (Berridge & Robinson, 2016). For example, a food photograph may activate perceptual and evaluative circuits, but if it predicts nothing during the task, it may not generate the anticipatory motivational state that mesolimbic dopamine encodes when cues reliably precede reward (Berridge & Robinson, 2003). Without actual juice delivery, the paradigm cannot dissociate wanting, the anticipatory signal elicited by the cue, from liking, the hedonic evaluation of the outcome itself, a distinction central to contemporary reward theories (Berridge & Kringelbach, 2015) This matters clinically: dysregulated incentive salience, where cue-triggered wanting drives behavior despite diminished liking or need, has been implicated in compulsive eating, obesity, and substance use disorders (Castro & Berridge, 2017; Morales & Berridge, 2020; Robinson & Berridge, 2025).

Gustatory neuroimaging studies address this gap by pairing visual cues with intra-oral liquid delivery during fMRI. This approach has identified anterior insula, orbitofrontal cortex, and ventral striatum as key regions for taste processing and outcome valuation (O’Doherty et al., 2002; Stice et al., 2008), with nucleus accumbens tracking reward anticipation in proportion to cue-outcome contingency (Knutson et al., 2001). Pavlovian conditioning paradigms have refined this picture: when arbitrary shapes are repeatedly paired with sweet liquid, the cues themselves come to activate reward-related regions including anterior insula, anterior cingulate, and orbitofrontal cortex, and cue-evoked motivated attention scales with the strength of the learned association (Cofresí & Aponte Zabala, 2026). These paradigms, however, achieve experimental control at the cost of ecological richness. Their cues are affectively neutral (typically geometric shapes), and therefore do not engage the natural selective attention systems that emotionally significant pictures recruit (Bradley, 2009; Versace et al., 2017). Picture-viewing and gustatory paradigms thus offer complementary strengths and weaknesses: the former engages naturalistic motivational salience but cannot deliver outcomes, while the latter delivers outcomes but uses cues stripped of motivational significance. It therefore remains unclear whether food cues embedded in a motivationally rich visual environment recruit striatal wanting signals beyond what their perceptual properties predict.

The present study integrates these two paradigms within a single fMRI session. We presented emotionally relevant images (erotica, mutilations, neutral scenes) and food photographs that either predicted an opportunity to receive sweet fruit juice (Food+) or did not (Food−). Because Food+ and Food− images were drawn from the same pool, any neural differences between conditions would reflect the learned outcome association rather than visual content. On each Food+ trial, participants indicated in real time whether they wanted the juice, enabling a within-subject dissociation of anticipatory signals during the cue period from consummatory signals during the outcome period.

We tested three predictions. First, we predicted that, relative to visually matched Food− cues, Food+ cues would recruit mesolimbic reward circuits, particularly the nucleus accumbens, reflecting a striatal wanting signal driven by learned contingency rather than perceptual or affective properties of the food image. Second, we predicted that anticipatory and consummatory signals would show dissociable neural patterns, with striatal regions dominating during the cue period and vmPFC during the outcome period, consistent with the wanting-liking distinction (Berridge & Kringelbach, 2015). Finally, we used representational similarity analysis to test which organizational principle best accounted for the multivariate structure of the neural response across our region-of-interest network: outcome availability, emotional arousal, affective valence, or visual category membership.

## Methods

### Participants

Twenty-three volunteers (14 female; mean age = 38.9 years, SD = 11.5, range: 25–62; mean BMI = 35.5, SD = 4.3) participated in the study. The sample was racially diverse (13 Black or African American, 7 White, 3 other; 6 Hispanic/Latino). Eighteen participants were right-handed. All participants provided written informed consent, and study procedures were approved by the University of Texas MD Anderson Cancer Center Institutional Review Board..

### Experimental paradigm

Participants completed six functional runs of a visuo-gustatory instructed prediction task, preceded by a short practice run to familiarize them with the trial structure and the juice delivery system. Stimuli were presented with E-Prime 2.0 (PST Inc., Pittsburgh, PA) and displayed via an MRI-compatible projection system; responses were collected with a Cedrus Lumina fiber-optic four-button response box. Participants were instructed to abstain from eating prior to the scanning session; if they reported being more than slightly hungry, a snack was provided before the session began.

On each trial, an image was presented for 2,000 ms followed by a variable inter-trial interval of 8, 10, or 12 s (Figure 1A). Pleasant, unpleasant, and neutral images were drawn from the International Affective Picture System (Lang et al., 2008) and from a database used in previous work (Versace et al., 2019). The 30 mutilation and 30 neutral images were all IAPS stimuli. Of the 30 erotica images, 22 were IAPS stimuli and 8 were in-house photographs matched in content and arousal. Of the 60 food images, 6 were IAPS stimuli and 54 were in-house photographs of palatable foods. The IAPS codes used were: neutral — 2037, 2039, 2102, 2107, 2190, 2191, 2210, 2273, 2305, 2359, 2374, 2377, 2383, 2393, 2396, 2397, 2411, 2435, 2441, 2500, 2511, 2512, 2575, 2594, 2595, 2620, 2630, 2635, 7550, 9070; mutilations — 3000, 3001, 3015, 3030, 3051, 3053, 3060, 3064, 3068, 3069, 3071, 3080, 3100, 3103, 3110, 3120, 3130, 3140, 3150, 3170, 3180, 3181, 3211, 3213, 3225, 3261, 3400, 6021, 9253, 9265; erotica — 4604, 4611, 4647, 4650, 4653, 4658, 4659, 4660, 4666, 4668, 4669, 4676, 4680, 4687, 4690, 4691, 4693, 4694, 4695, 4696, 4698, 4800; food — 7330, 7390, 7405, 7410, 7430, 7470. Pleasant images consisted of erotica and unpleasant images consisted of mutilation scenes; these categories were selected because they are highly arousing and reliably engage motivational circuits (Sabatinelli et al., 2005). Throughout the paper, “Pleasant” and “Unpleasant” refer to these specific stimulus categories. Food+ and Food− images were photographs of highly palatable foods drawn from the same pool. Each food photograph was presented within either a red or a yellow frame, and for each participant one color signaled Food+ trials while the other signaled Food− trials, with the color-to-condition mapping counterbalanced across participants. Photograph assignment to Food+ versus Food− was likewise counterbalanced across participants, so any neural difference between Food+ and Food− could not be attributed to visual content.

**Figure 1.**
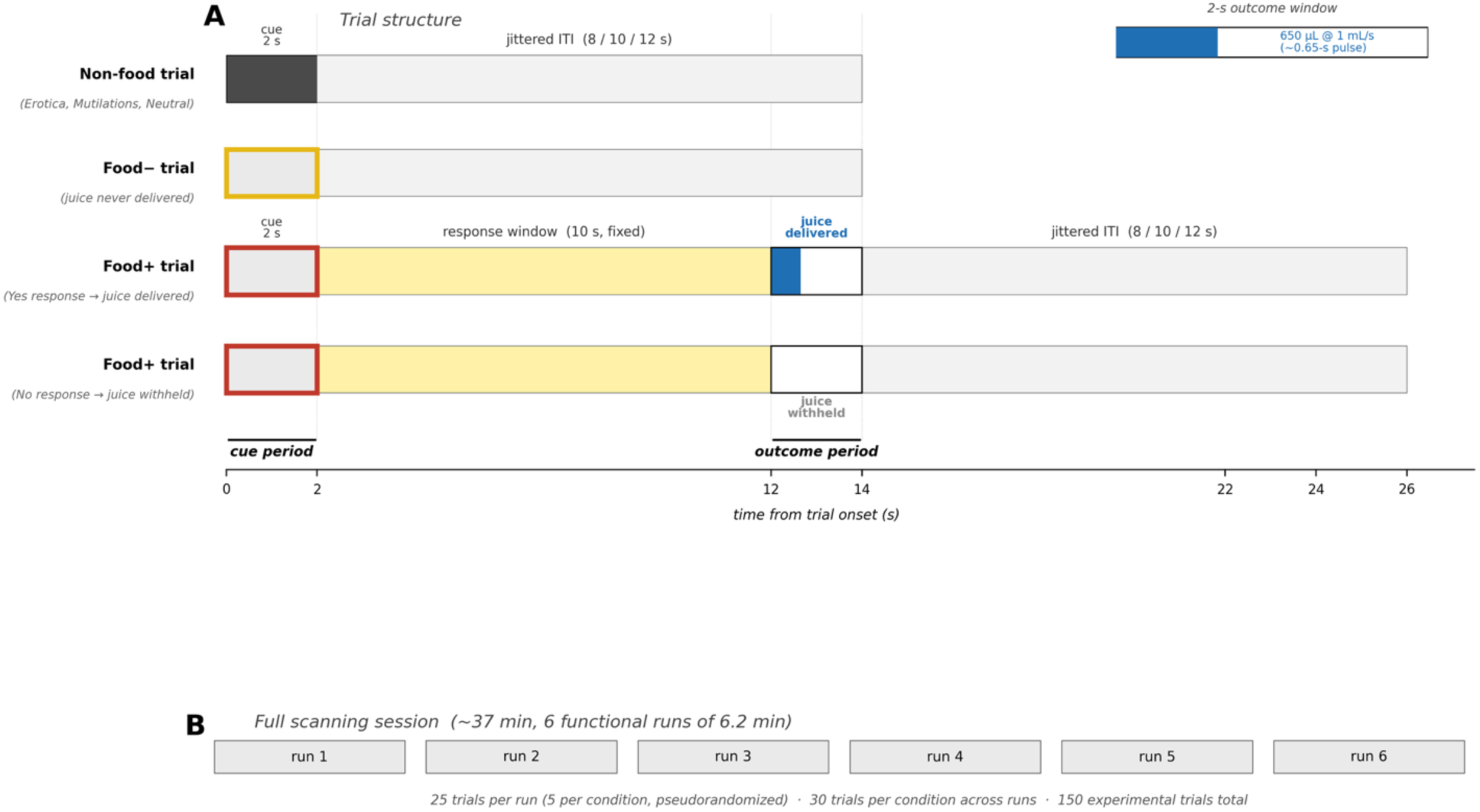
Experimental paradigm and timing. (A) Trial-level structure for the four trial types. Food images appeared within a colored frame: red for Food+ (juice-eligible) and yellow for Food− (juice-ineligible) in the schematic shown, with the color-to-condition mapping counterbalanced across participants. Erotica, Mutilations, and Neutral images had no frame. Non-food trials (Erotica, Mutilations, Neutral, Food−) consisted of a 2-s image followed by a jittered inter-trial interval of 8, 10, or 12 s. Food+ trials began with a 2-s Food+ image followed by a fixed 10-s response window during which participants indicated by button press whether they wanted to receive the juice. The 2-s outcome window started at 12 s after trial onset: on Yes-response trials, 650 µL of orange juice was delivered through an MRI-compatible syringe pump at 1 mL/s (∼0.65-s pulse, inset); on No-response trials, no juice was delivered, but the trial timing was otherwise identical. A second jittered interval (8, 10, or 12 s) followed the outcome window. We refer to the 2-s image-presentation window as the cue period and to the 2-s juice-delivery window as the outcome period throughout. (B) Full scanning session, comprising six functional runs of 6.2 min each (∼37 min of functional scanning in total). Each run contained 25 trials (5 per condition), pseudorandomized within runs with the constraint that no two consecutive trials belonged to the same condition, for a total of 30 trials per condition (150 experimental trials in total).

Food+ trials, and only Food+ trials, included an opportunity to receive juice. After the Food+ image, participants had a fixed 10-s response window (5 volumes) to indicate by button press whether they wanted to receive juice on that trial (Yes/No, with left/right button assignment counterbalanced across participants). At the end of this response window, juice was delivered following Yes responses and withheld following No responses; if no response was recorded, juice was delivered by default. The outcome was followed by a variable interval of 8, 10, or 12 s before the next trial. On Yes trials, approximately 650 μl of room-temperature orange juice was delivered through vinyl tubing to a pacifier-style mouthpiece held between the participant’s lips. Delivery was controlled by a syringe pump, which advanced the syringe plunger and pushed juice through the tubing. On No trials, no juice was delivered but timing was otherwise identical, allowing cue-evoked anticipatory signals to be dissociated from outcome-related consummatory signals at the contrast level.

Each run lasted 6.2 minutes and comprised 186 volumes (including 3 initial dummy volumes) and began with a checkerboard trial that was excluded from analyses (Figure 1B). Each run included 25 trials, five per condition (Erotica, Mutilations, Neutral, Food+, Food−), pseudorandomized within runs with the constraint that no two trials from the same category appeared consecutively. Across the six runs, each participant saw 30 trials per condition (150 experimental trials in total).

Perceived satiety was assessed before and after the scanning session using the Satiety Labeled Intensity Magnitude scale (SLIM) (Cardello et al., 2005). After the fMRI session, participants rated each picture for valence and arousal, using the 9-point scale Self-Assessment Manikin (SAM) (Bradley & Lang, 1994). Valence and arousal ratings were analyzed with separate one-way repeated-measures ANOVAs, with picture category (food, erotica, neutral scenes, mutilations) as the within-subject factor. Greenhouse–Geisser correction was applied where sphericity was violated. Pairwise post-hoc comparisons were performed using Bonferroni corrected paired-samples t-tests.

### fMRI acquisition

Data were collected at MD Anderson Cancer Center on a Siemens Prisma 3T scanner with a 20-channel head/neck coil. Functional volumes were acquired using a T2*-weighted echo planar imaging sequence (ep2d_bold) with TR = 2,000 ms, TE = 30 ms, flip angle = 90°, voxel size = 3.0 × 3.0 × 3.0 mm (0.51 mm inter-slice gap), 37 axial slices, and GRAPPA acceleration factor of 2. Each of the six functional runs comprised 183 volumes (366 s). A high-resolution T1-weighted anatomical scan (MPRAGE) was also acquired (TR = 2,200 ms, TE = 2.45 ms, TI = 900 ms, flip angle = 8°, voxel size = 0.9375 × 0.9375 × 1.0 mm, 176 slices, GRAPPA factor = 2).

### fMRI preprocessing

Preprocessing was performed using AFNI (Cox, 1996) via the `afni_proc.py` pipeline (Reynolds et al., 2024). Each participant’s T1-weighted anatomical image was skull-stripped and nonlinearly warped to the MNI152 2009 asymmetric template with @SSwarper. Functional runs then underwent temporal despiking (`3dDespike`), slice-timing correction for an interleaved ascending acquisition, rigid-body motion correction with the volume of lowest outlier fraction as the alignment target, affine coregistration of the EPI to the anatomical image (lpc+ZZ cost function, `giant_move` initialization), and nonlinear normalization to MNI space. Motion correction, EPI-to-anatomical alignment, and the nonlinear warp were concatenated and applied in a single interpolation step to minimize spatial smoothing. Data were then smoothed with a 6 mm FWHM Gaussian kernel and scaled to percent signal change relative to each run’s mean. Volumes were censored from subsequent analyses if the frame-wise Euclidean norm of the motion derivative exceeded 0.3 mm or if the fraction of outlier voxels exceeded 0.1. A subject-specific grey-matter mask, derived from the T1-weighted image via CAT12 (Gaser et al., 2024) segmentation and resampled to the functional resolution, was applied to the output statistical maps to restrict analyses to grey matter.

### First-level analysis

Individual subject data were analyzed using 3dDeconvolve in AFNI. The general linear model included regressors for the checkerboard trial at the start of each run, for Erotica, Mutilations, Neutral, and Food− cues, Food+ cues and the outcome period. All regressors were modeled as 2-s boxcars convolved with AFNI’s canonical hemodynamic response (BLOCK(2,1) basis function). Food+ cue onsets were modeled with amplitude modulation (AM2): a single regressor capturing the mean Food+ response was accompanied by a parametric modulator coding the participant’s subsequent choice (Yes / No, mean-centered) and a linear modulator capturing slow drift across the session. The outcome period was modeled analogously, with parametric modulators for delivery (Yes/No) and session drift. The choice modulator beta on the Food+ regressor therefore indexed the differential cue-period response on trials subsequently accepted versus declined; the delivery modulator beta on the juice regressor indexed the differential outcome-period response on trials with versus without juice receipt. Six head-motion parameters and their derivatives were included as nuisance regressors, and motion-censored TRs (frame-wise displacement > 0.3 mm or outlier fraction > 0.1) were excluded. This procedure yielded condition-specific beta maps for each participant.

Four participants accepted the juice on 100% of trials, contributing zero variance to the choice modulator. For these participants, Food+ and juice were modeled with plain BLOCK(2,1) regressors without amplitude modulation; their data contributed to all mean-activation contrasts (N = 23) but were excluded from the Yes vs. No contrasts (N = 19).

### ROI definition

Two sets of ROIs were defined for complementary purposes.

#### Functional ROIs

To validate the paradigm, we identified regions responsive to emotional content using a whole-brain voxel-wise contrast of Emotional (Erotica + Mutilations) versus Neutral. Four functional ROIs were defined from activation clusters in this contrast: dorsal and ventral visual cortex, medial prefrontal cortex (mPFC), inferior frontal gyrus (IFG), and amygdala. These regions were used exclusively for paradigm validation analyses.

#### A priori ROIs

Six bilateral ROIs were defined from independent anatomical atlases based on their theoretical relevance to food reward processing, interoception, and value-based decision making (Figure 3A). Subcortical ROIs were extracted from the Tian Subcortex Atlas Scale II (3T) (Tian et al., 2020): nucleus accumbens (NAc; combining shell and core subdivisions) and putamen (combining anterior and posterior subdivisions). Ventromedial prefrontal cortex (vmPFC) was defined as the Frontal Medial Cortex region of the Harvard-Oxford Cortical Atlas (probability threshold >25%). Three insula subdivisions were defined from a resting-state functional connectivity parcellation (Deen et al., 2011): dorsal anterior insula (dAI), ventral anterior insula (vAI), and posterior insula (PI). All ROIs were bilateral (left and right hemispheres collapsed) and resampled to the functional data space using nearest-neighbor interpolation.

For each participant and condition, mean beta values were extracted from each ROI.

### Statistical analysis

All ROI-level analyses were conducted in Python (version 3.14.0) using SciPy and NumPy.

#### Within-subject contrasts

For each ROI, condition-specific betas were tested against zero using one-sample t-tests. Paired comparisons between conditions (e.g., Erotica vs. Neutral, Food+ vs. Food−) were computed as within-subject difference scores and tested via one-sample t-tests on the difference maps. Yes versus No contrasts (Food+ Yes > No during the cue period; Juice Yes > No during the outcome period) were computed analogously for the subset of participants (N = 19) who made at least one Yes and one No choice across the experiment.

#### Multiple comparison correction

To control for multiple comparisons across ROIs, we applied the Benjamini-Hochberg false discovery rate (FDR) procedure within three theoretically motivated test families: (1) emotional validation comparisons across four functional ROIs, (2) food processing comparisons across six a priori ROIs, and (3) Yes/No contrasts across six a priori ROIs. All p-values reported from ROI analyses reflect FDR-corrected values within their respective family unless otherwise noted. Effect sizes are reported as Cohen’s d for paired comparisons.

#### Representational similarity analysis (RSA)

To characterize how the neural response differentiated the five experimental conditions across the full region-of-interest network, we performed a model-based representational similarity analysis (RSA) using the same condition-specific mean betas entered in the univariate ROI analyses reported above. For each participant, we assembled a 5 × 10 activation profile: five conditions (Neutral, Erotica, Mutilations, Food−, Food+) by ten ROIs. The ten ROIs comprised the four functional ROIs (visual cortex, mPFC, IFG, and amygdala) and the six a priori ROIs (NAc, putamen, vmPFC, dAI, vAI, and PI). From these profiles, we computed a 5 × 5 neural representational dissimilarity matrix (RDM) by taking the correlation distance (1 − Pearson r) between the 10-ROI profiles of each pair of conditions. A pair of conditions has a low dissimilarity if the 10-ROI network responds to them with similar profiles, and a high dissimilarity if their profiles differ. Importantly, the input to the RSA is the same set of subject-level condition betas plotted as univariate violin plots in the functional ROI and a priori ROI analyses. The Yes/No contrasts reported in the temporal dissociation analysis below were not used.

We compared each participant’s neural RDM to four theoretical model RDMs, each formalizing a distinct hypothesis about what organizes the neural response to the five conditions. A model RDM specifies the pairwise dissimilarity pattern expected if that hypothesis is correct. The Food Reward model tested whether outcome prediction organized the neural response: because Food+ was the only condition paired with imminent reward delivery, it should be representationally distinct from all other conditions, which should be mutually similar. The Arousal model tested whether motivational intensity organized the response: the two high-arousal emotional conditions (Erotica, Mutilations) should cluster together and be dissimilar from the three low-arousal conditions (Neutral, Food−, Food+). The Valence model tested whether affective valence organized the response: Erotica (pleasant) and Mutilations (unpleasant) should be maximally dissimilar, with Neutral and food conditions occupying intermediate positions. The Visual Category model tested whether perceptual category organized the response: stimulus categories were predicted to be mutually dissimilar, with the exception of Food+ and Food−, which were predicted to cluster together because both consisted of food photographs.

For each participant, the neural RDM was correlated with each model RDM using Spearman’s ρ on the vectorized lower triangles. Group-level significance for each model was assessed via 10,000 permutations in which the five condition labels were randomly shuffled within each participant before recomputing the neural RDMs and their correlation with each model. Pairwise comparisons between models were conducted using paired t-tests on subject-level ρ values. A noise ceiling was estimated using both upper-bound (correlation of each subject’s neural RDM with the group-mean neural RDM) and lower-bound (correlation with the leave-one-out mean neural RDM) approaches (Nili et al., 2014).

#### Exploratory satiety analyses

To examine whether pre-scan perceived satiety modulated food-cue-related brain responses, we correlated SLIM scores with behavioral juice acceptance rate (%Yes) and with ROI-level betas for three food-specific contrasts (Food+ minus Neutral, Food− minus Neutral, Food+ minus Food−) and the two Yes/No contrasts (Food+ Yes > No, Juice Yes > No). Both correlations were computed as zero-order Pearson r and as partial correlations controlling for BMI and age. Because SLIM scores were available for 22 of the 23 participants (18 of the 19 with variable Yes/No behavior), these analyses used the corresponding subsamples. Correction for multiple comparisons was applied using the Benjamini-Hochberg FDR procedure within two families: food-specific contrasts (3 contrasts × 10 ROIs = 30 tests) and Yes/No contrasts (2 contrasts × 10 ROIs = 20 tests).

## Results

### Affective ratings

Affective ratings, used as a manipulation check, confirmed the expected emotional differentiation across picture categories. For valence, the repeated-measures ANOVA revealed a large main effect of category, F(3, 49.27) = 111.05, p < .001, η²p = .84 (Greenhouse–Geisser corrected, ε = .78). Post-hoc comparisons showed that mutilations received the lowest valence ratings (M = 1.81, SEM = 0.20) and were rated as significantly more negative than all other categories (all p < .001, d > 2.6). Food images (M = 6.55, SEM = 0.24) and erotica (M = 6.18, SEM = 0.28) were both rated as pleasant and did not differ from each other (p = 1.00), but both were rated significantly more positive than neutral scenes (M = 5.13, SEM = 0.14; p < .001 and p = .003, respectively).

For arousal, the ANOVA also revealed a significant main effect of category, F(3, 50.15) = 17.24, p < .001, η²p = .45 (ε = .80). Mutilations were rated as the most arousing category (M = 6.48, SEM = 0.41), significantly higher than erotica (M = 4.93, SEM = 0.42; p = .034, d = 0.66), food images (M = 4.32, SEM = 0.40; p = .005, d = 0.82), and neutral scenes (M = 3.16, SEM = 0.32; p < .001, d = 1.39). Erotica and food images did not differ in arousal (p = 1.00), but both were rated as significantly more arousing than neutral scenes (erotica: p < .001, d = 1.04; food: p = .014, d = 0.74).

Within the food category, ratings did not differ between Food+ and Food− images. Valence was virtually identical for the two conditions (Food+ M = 6.57, SEM = 0.25; Food− M = 6.52, SEM = 0.24; t(21) = 0.46, p = .650, d = 0.10), and arousal was also comparable (Food+ M = 4.43, SEM = 0.43; Food− M = 4.18, SEM = 0.37; t(21) = 1.39, p = .180, d = 0.30).

### Behavioral results on juice acceptance

Participants showed substantial individual variability in juice acceptance across the experiment. The overall acceptance rate averaged 56.5% (SD = 32.6%, range: 3.3–100%), reflecting heterogeneity in the motivational value attributed to the gustatory reward across individuals and across the session (Figure 2A, B). Within-subject acceptance rates were stable enough to be meaningful yet variable enough to support neural dissociations: the majority of participants neither uniformly accepted nor uniformly rejected the juice, indicating that choices reflected fluctuating internal states rather than habitual responding. Four participants accepted the juice on every trial and were therefore excluded from the Yes vs. No neural contrasts (yielding N = 19 for those analyses); they were retained for all cross-condition comparisons (N = 23).

**Figure 2.**
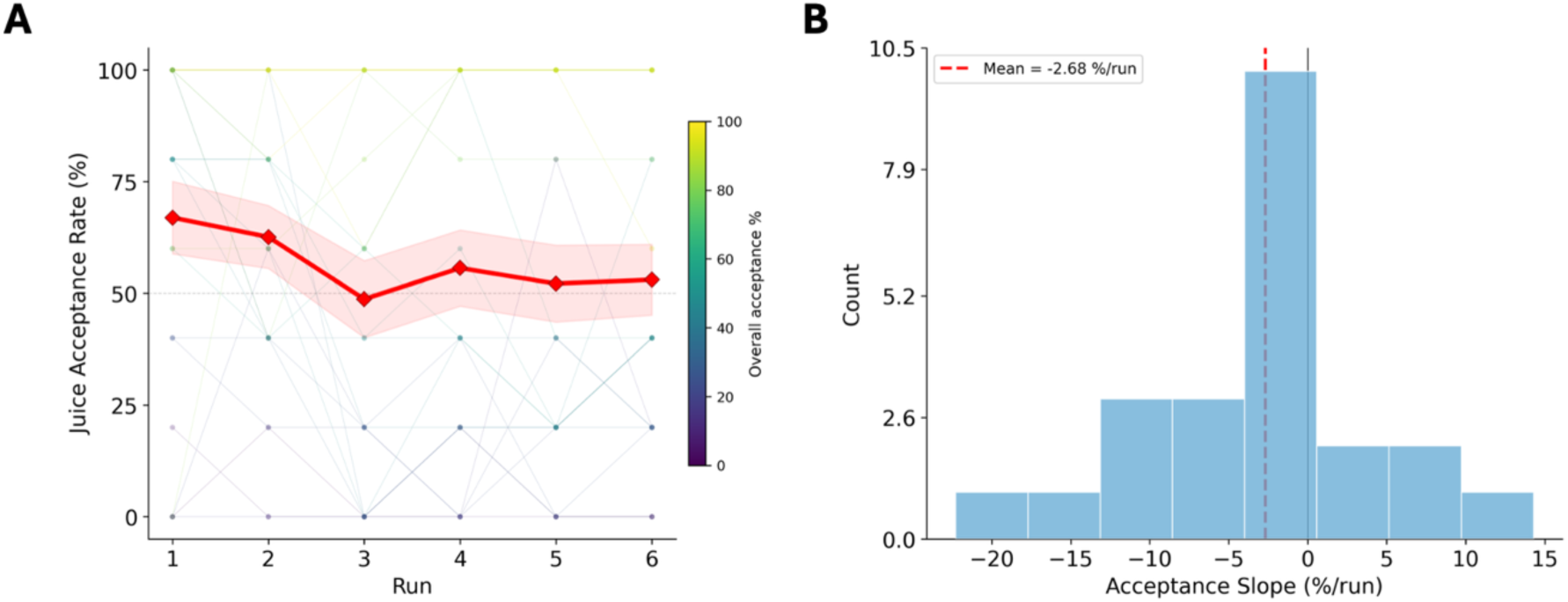
Behavioral characterization of juice acceptance. (A) Individual subject trajectories of juice acceptance rate (% Yes choices on Food+ trials) across the six scanning runs. Each line represents one participant, colored by overall acceptance rate (viridis colormap). The red line with diamond markers shows the group mean (±1 SEM shading). The dashed grey line marks 50% acceptance. (B) Distribution of within-subject linear slopes of acceptance rate across runs. The red dashed line marks the group mean slope (−2.68%/run). The wide distribution of slopes reflects substantial individual variability in how acceptance changed across the session, with 13 of 23 participants showing declining and 10 showing stable or increasing acceptance.

### Paradigm validation: Emotional processing in functional ROIs

To confirm that the paradigm evoked robust emotional processing, we examined whole-brain voxel-wise activation for the Emotional > Neutral contrast and extracted condition-specific betas from four functional regions defined by this contrast (Figure 3).

**Figure 3.**
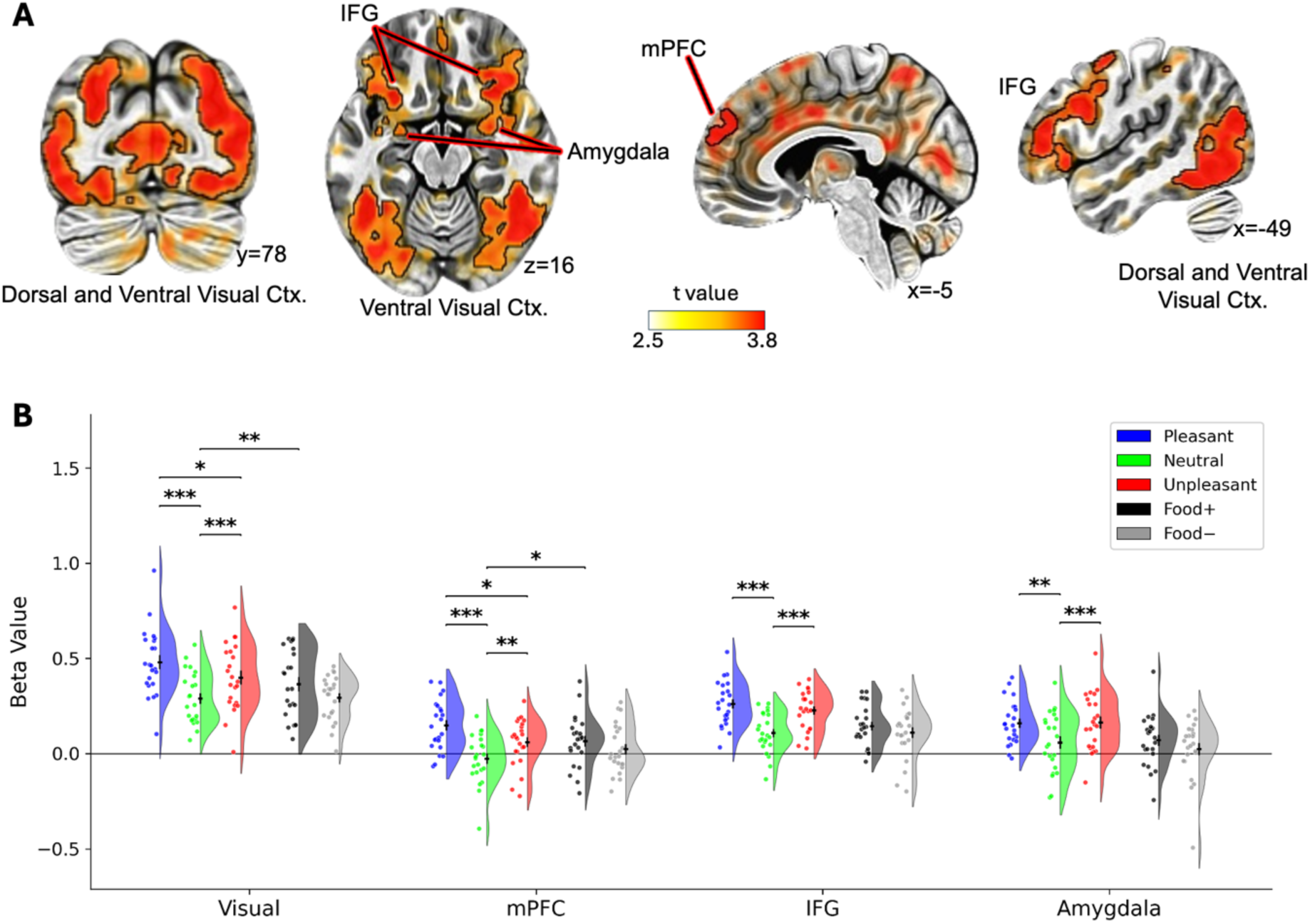
Paradigm validation: emotional and food cue processing in functional ROIs. (A) Whole-brain voxel-wise activation maps for the Emotional > Neutral contrast, displayed on a standard MNI template. Anatomical labels indicate the four functional ROIs defined from this contrast: dorsal and ventral visual cortex, medial prefrontal cortex (mPFC), inferior frontal gyrus (IFG), and amygdala. Coordinates are +LPI in MNI space. (B) Mean beta values (±1 SEM) extracted from each of the four functional ROIs for all five experimental conditions: Pleasant (Erotica, blue), Neutral (green), Unpleasant (Mutilations, red), Food+ (black), and Food− (grey). Brackets with asterisks denote significant pairwise comparisons (paired t-tests, FDR-corrected within family). ***p < .001, **p < .01, *p < .05. Coordinates

The whole-brain analysis revealed extensive bilateral activation of visual cortex, mPFC, IFG, and amygdala for emotional relative to neutral images (Figure 3A). Subsequent analyses confirmed this pattern (Figure 3B): in all four functional regions, both Erotica and Mutilations elicited significantly greater activation than Neutral (all pFDR < .005; Figure 3B). Effect sizes were large, ranging from d = 0.66 (Amygdala, Erotica vs. Neutral) to d = 1.85 (Visual, Erotica vs. Neutral).

Critically, Food+ images also engaged a subset of these regions. In visual cortex, Food+ elicited significantly greater activation than Neutral (pFDR = .009, d = 0.66), and in mPFC, Food+ exceeded Neutral (pFDR = .041, d = 0.54; Figure 3B). However, Food+ did not significantly exceed Neutral in IFG or amygdala (both pFDR > .15), and Food− did not differ from Neutral in any functional region (all pFDR > .50). This differential pattern indicates that Food+ images, which predict actual reward delivery, engage visual cortex and mPFC similarly to emotional stimuli, whereas Food− images, which have no outcome association do not.

### Food cue processing in a priori ROIs

To characterize neural responses to food cues across reward, valuation, and interoceptive circuits, we examined the six a priori ROIs (Figure 4A).

**Figure 4.**
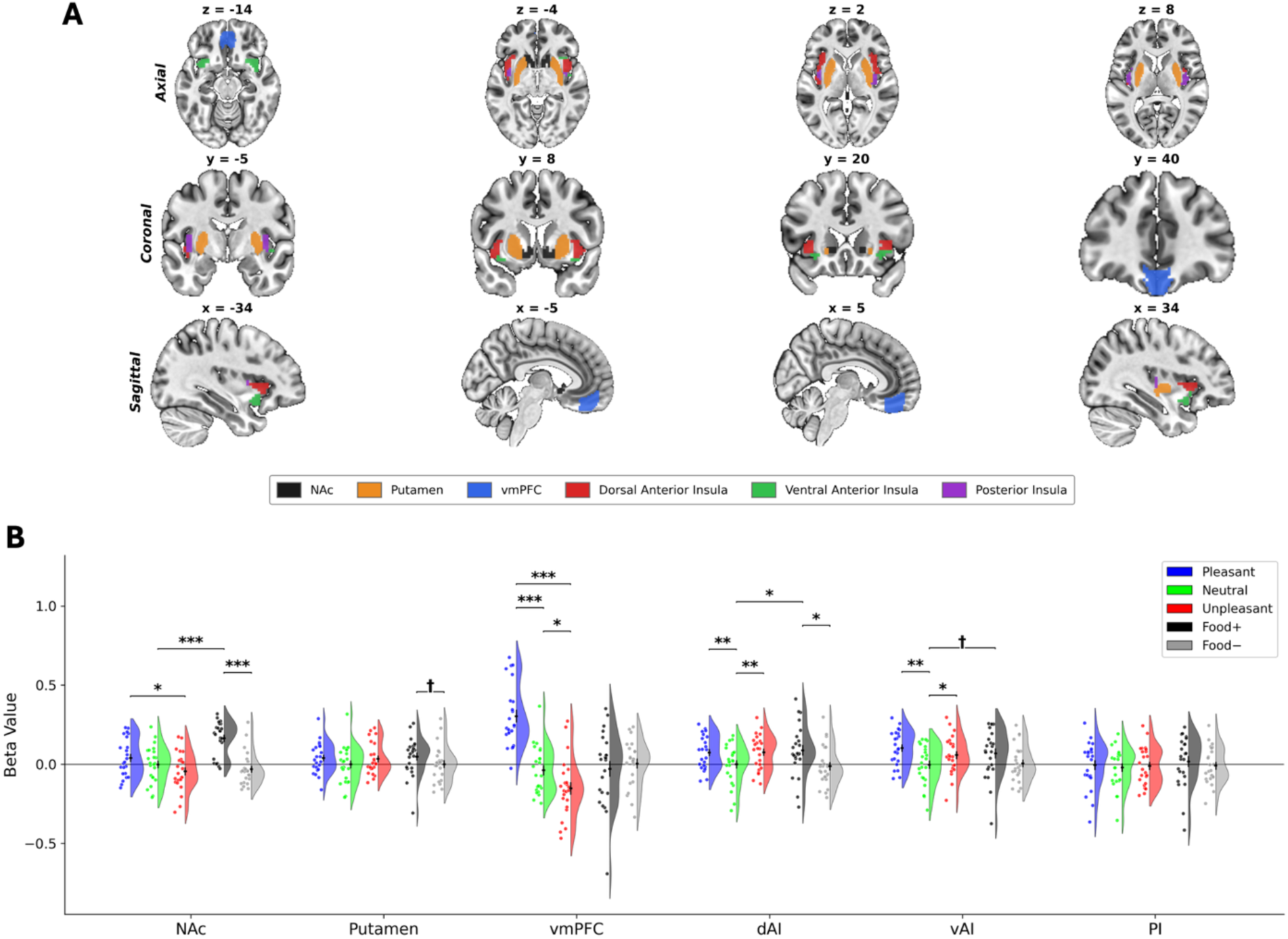
Food cue processing in a priori ROIs. (A) Anatomical location of the six bilateral a priori ROIs displayed on a standard MNI template in axial, coronal, and sagittal views. NAc and putamen were defined from the Tian Subcortex Atlas (Scale II, 3T). vmPFC was defined from the Harvard-Oxford Cortical Atlas (Frontal Medial Cortex). Dorsal anterior insula (dAI), ventral anterior insula (vAI), and posterior insula (PI) were defined from resting-state functional connectivity parcellation (Deen et al., 2011). (B) Mean beta values (±1 SEM) for emotional conditions (left panel) and food conditions (right panel) across the six a priori ROIs. Brackets with asterisks denote significant pairwise comparisons (paired t-tests; *p < .001, p < .01, *p < .05, †p < .1).

#### Emotional responses

In the Emotional conditions panel (Figure 4B, left), vmPFC showed a clear valence dissociation: Erotica elicited consistent positive activation (d = 1.55, p < .001) while Mutilations produced significant negative deflection (d = −0.81, p < .001), with the Erotica vs. Mutilations difference being the largest observed in any ROI (d = 2.46, p < .001). Both dAI and vAI responded significantly to emotional stimuli relative to Neutral (dAI: Erotica pFDR = .005, Mutilations pFDR = .014; vAI: Erotica pFDR = .003, Mutilations pFDR = .035). NAc, putamen, and PI showed no significant emotional responses (all pFDR > .10).

#### Food-cue responses

The Food conditions (Figure 4B) revealed a striking pattern. NAc showed a large selective response to Food+ (d = 1.51, p < .001), with no response to any other condition, and a highly significant Food+ vs. Food− difference (d = 1.28, pFDR < .001). This Food+ signal in NAc was significantly larger than both Erotica (p < .001) and Mutilations (p < .001), indicating that NAc is not tracking emotional arousal, rather the anticipation of a rewarding outcome. In dAI, Food+ significantly exceeded both Neutral (p= .034) and Food− (p= .036), though these did not survive FDR correction (pFDR = .16). vAI showed a trending Food+ vs. Neutral effect (p= .086, pFDR = .29). Putamen, vmPFC, and PI showed no significant food condition effects (all pFDR > .15).

### Representational similarity analysis across the ROI network

The univariate analyses reported above examined, one region at a time, how each experimental condition activated each ROI. We next asked a complementary multivariate question: across the full 10-ROI network, which conditions elicited similar vs. distinct patterns of representation? To answer this question, we used the same condition-specific mean betas shown in Figures 3 and 4 to perform a model-based representational similarity analysis (RSA; Figure 5).

**Figure 5.**
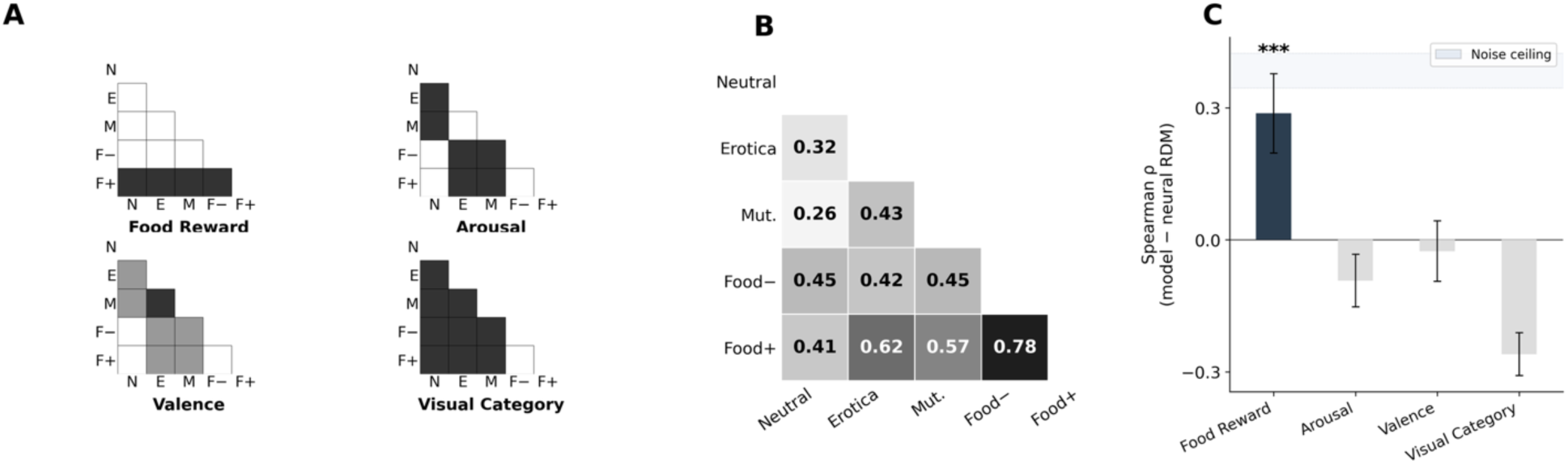
Model-based representational similarity analysis. (A) Predicted representational dissimilarity matrices (RDMs) for four competing theoretical models. Each matrix specifies the predicted pairwise dissimilarity between conditions (N = Neutral, E = Erotica, M = Mutilations, F− = Food−, F+ = Food+), where dark cells indicate high predicted dissimilarity and light cells indicate low predicted dissimilarity. (B) Observed group-mean neural RDM, computed as the average correlation distance (1 − Pearson r) between 10-ROI activation profiles for each condition pair (N = 23). The Food+ row shows the highest dissimilarity values, particularly with Food− (0.78), despite these conditions sharing identical visual content. (C) Model fit to the neural data. For each participant, the neural RDM was correlated (Spearman ρ) with each model RDM. Bars show group-mean correlations (±1 SEM). Significance was assessed via permutation testing (10,000 iterations). Only the Food Reward model significantly predicted the neural data (p < .001). The shaded band indicates the noise ceiling estimated from upper and lower bounds.

We tested four theoretical models (Figure 5A), each specifying a different account for how the five conditions should be organized in the neural representational space. <u>The Food Reward model</u> predicted that Food+ would be isolated because it is the only reward-predictive cue. <u>The Arousal</u> <u>model</u> predicted that Erotica and Mutilations would cluster together as the high-arousal conditions. <u>The Valence model</u> predicted that Erotica and Mutilations would be maximally dissimilar as the two affective poles, with the low-arousal conditions at intermediate distances. <u>The Visual Category</u> <u>model</u> predicted that each visual category would be represented as distinct from the others, except that Food+ and Food− would cluster together as food images.

The group-mean observed neural RDM (Figure 5B) revealed a striking asymmetry in how the 10-ROI network represented the five conditions. Food+ was highly dissimilar from every other condition, and the largest dissimilarity in the matrix was between Food+ and Food− (0.78). This finding is notable because Food+ and Food− were drawn from the same set of food photographs and differed only in whether they predicted an imminent reward delivery. By comparison, the dissimilarity between Erotica and Mutilations, two high-arousal conditions at opposite ends of the valence spectrum, was 0.43. Thus, within this ROI network, the representational separation associated with actual reward delivery prediction exceeded the separation between the most pleasant and most unpleasant images in the task.

Permutation testing confirmed that only the Food Reward model significantly predicted the observed neural data (ρ = 0.29, p < .001; Figure 5C). The Arousal (ρ = −0.09, p = .84), Valence (ρ = −0.02, p = .56), and Visual Category (ρ = −0.26, p > .99) models all failed to reach significance. Pairwise comparisons confirmed that the Food Reward model fit significantly better than each of the three alternatives (all p < .001). Across the 10-ROI network, outcome prediction, not emotional arousal, affective valence, or visual category membership, was the dominant organizing principle of the multi-region neural response.

### Temporal dissociation of anticipatory and consummatory signals

To dissociate neural activity during cue anticipation from activity during the outcome period, corresponding conceptually to anticipatory wanting and consummatory liking, we computed two within-subject difference contrasts across all 10 ROIs in the 19 participants who made both Yes and No choices (Figure 6).

**Figure 6.**
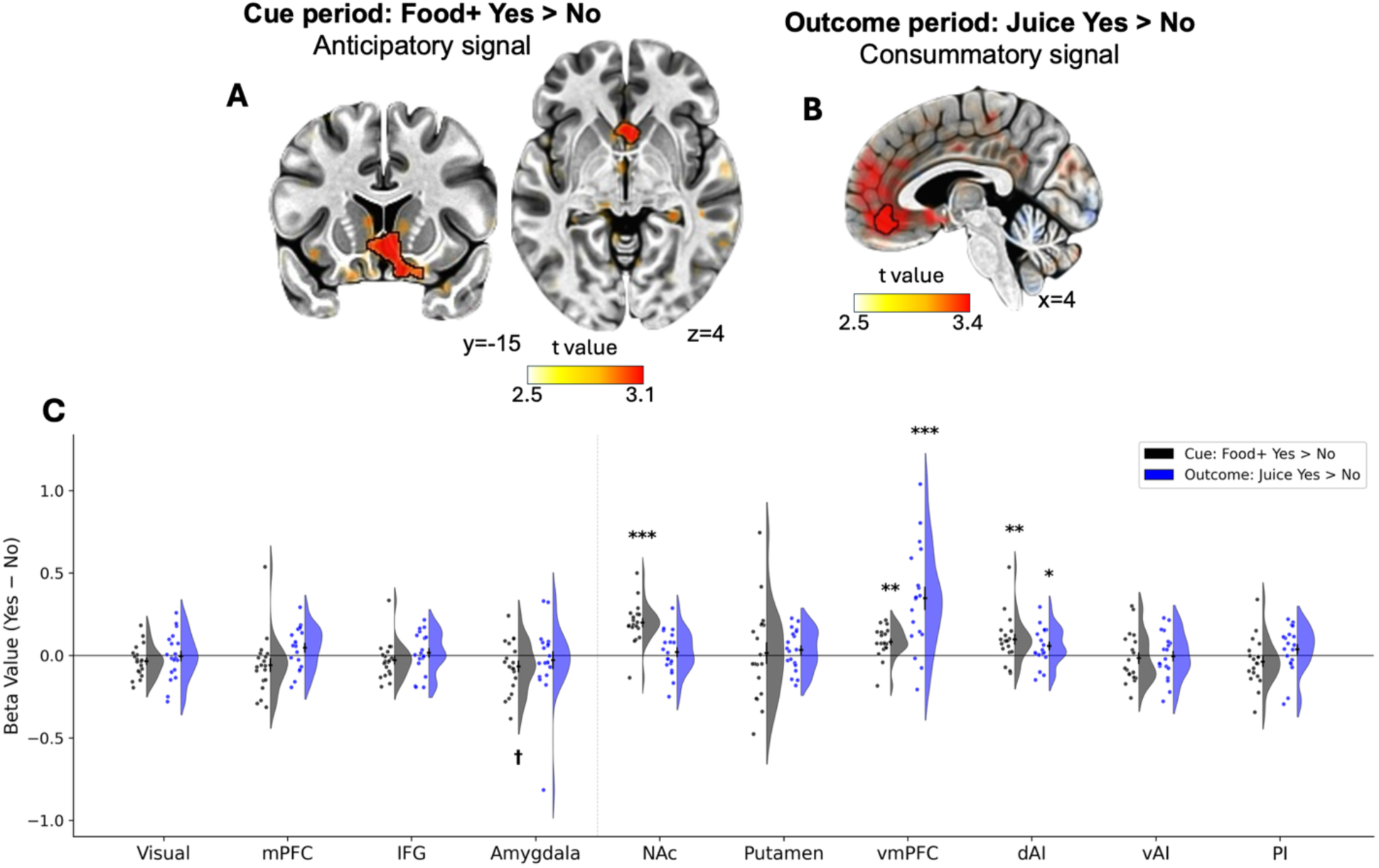
Temporal dissociation of anticipatory and consummatory signals. (A) Whole-brain voxel-wise map for the Food+ Yes > No contrast during the cue period (anticipatory signal), displayed on a standard MNI template. The highlighted cluster shows ventral striatum activation. (B) Whole-brain voxel-wise map for the Juice Yes > No contrast during the outcome period (consummatory signal), displayed on a sagittal view of the medial wall. The highlighted cluster shows ventromedial prefrontal cortex activation. Coordinates are +LPI in MNI space. (C) Mean beta values (±1 SEM) for the Food+ Yes > No contrast (black bars) and Juice Yes > No contrast (blue bars) across all 10 ROIs (N = 19). The dashed vertical line separates functional ROIs, derived from the whole-brain Emotional > Neutral contrast (left), from a priori anatomical ROIs (right). All values shown are within-subject Yes minus No differences; regions that respond similarly on Yes and No trials will show values near zero even if their absolute activation to Food+ or juice is high. Asterisks denote significance of one-sample t-tests against zero (FDR-corrected within the Yes/No contrast family). During cue processing, anticipatory signals favoring upcoming Yes choices are concentrated in NAc, vmPFC, and dAI. During outcome processing, consummatory signals are dominated by vmPFC, with dAI also participating. ***p < .001, **p < .01, *p < .05.

The cue-period contrast, Food+ Yes > No, compared Food+ trials that were later accepted, leading to juice delivery, with Food+ trials that were later declined, leading to no juice delivery. Because both trial types involved the same food images, this contrast isolated the component of cue-period activity associated with the subsequent decision to accept the juice.

The outcome-period contrast, Juice Yes > No, compared accepted Food+ trials, in which the juice was delivered, with declined Food+ trials in which no juice was delivered. This contrast isolated the component of outcome-period activity associated with actual juice receipt.

In both cases, activation on trials in which the participant declined the juice (“No”) was subtracted from activation on trials in which the participant accepted it (“Yes”), thereby removing stimulus-driven activity shared across trial types.

As illustrated in Figure 6, during the cue period, Food+ Yes > No was significant in NAc (d = 1.59, pFDR < .001), vmPFC (d = 0.90, pFDR = .003), and dAI (d = 0.69, pFDR = .020). No other ROI showed a significant cue-period effect (all pFDR > .15). During the outcome period, Juice Yes > No was significant in vmPFC (d = 1.11, pFDR < .001) and dAI (d = 0.52, pFDR = .072, trending). No other ROI showed a significant outcome-period effect (all pFDR > .25).

#### Exploratory associations with pre-scan satiety

Pre-scan perceived satiety (SLIM score; mean = −3.6, SD = 33.5, range: −70.8 to 73.6) was not associated with juice acceptance rate across subjects (r = −0.21, p = .36), suggesting that the behavioral variability driving the Yes/No contrasts reflects fluctuating motivational state rather than current hunger. SLIM scores were uncorrelated with both BMI (r = −0.18, p = .43) and age (r = +0.09, p = .71).

In exploratory partial correlations controlling for BMI and age, SLIM scores were positively associated with the Food+ minus Neutral contrast in dorsal anterior insula (r = +0.45, p = .034) and ventral anterior insula (r = 0.44, p = .041), and with the amygdala Juice Yes > No contrast at outcome (r = +0.60, p = .009). No association with SLIM was observed for NAc, vmPFC, putamen, or any other ROI in any contrast (all p > .13). The direction of these associations indicates that individuals reporting greater satiety showed stronger anterior insula differentiation of Food+ from neutral scenes and stronger amygdala differentiation of reward delivery from withholding.

However, none of these associations survived FDR correction within their respective test families (anterior insula: pFDR = .62; amygdala: pFDR = .17), and they should therefore be considered exploratory observations pending replication in larger samples.

## Discussion

The present study bridged two traditions in human reward neuroimaging: free viewing of emotionally significant pictures and visuo-gustatory instructed prediction with real reward delivery, to test whether food cues predicting actual juice delivery recruit striatal wanting signals beyond those evoked by visually matched, non-predictive food cues. Three converging results indicate that instructed outcome prediction dominated the neural response to food cues in this paradigm. NAc responded selectively to Food+ cues, with greater activation than to both high-arousing pleasant (erotica) and unpleasant (mutilation) images. Cue-period anticipatory signals and outcome-period consummatory signals were anatomically dissociable, with NAc activity dominant during anticipation and vmPFC activity dominant after the outcome period. Representational similarity analysis further showed that outcome prediction, rather than arousal, valence, or visual category, best accounted for the multi-region pattern of neural response. Together, these findings show that, in this paradigm, whether a visual stimulus engages mesolimbic reward circuitry depends more on what the stimulus predicts than on what it depicts.

### Outcome prediction drives striatal wanting signals

The selectivity of the NAc response to Food+ cues was the central finding of the study. NAc activation was substantially larger for Food+ than for Food−, and exceeded NAc responses to erotica, a condition typically used as a benchmark of appetitive mesolimbic engagement (Demos et al., 2012; Sabatinelli et al., 2007). This differential pattern is unlikely to be explained by visual content, because Food+ and Food− were drawn from the same pool of food photographs. It is also unlikely to be explained by arousal: in arousal-sensitive regions of our network, including visual cortex and amygdala, Food+ activation was substantially lower than activation to erotica, indicating that Food+ images are not more arousing than erotica by standard neural markers, consistent with prior findings that food pictures, despite being rated as arousing, do not reliably evoke neuroaffective responses comparable to erotica or mutilations in passive-viewing contexts (Versace et al., 2023). Yet in NAc, Food+ exceeded erotica. The arousal account cannot reconcile these patterns.

Instead, the critical difference between the two food conditions was whether the cue predicted immediate reward delivery. This pattern is consistent with the established role of NAc in anticipatory reward and incentive salience attribution (Knutson et al., 2001; Robinson & Berridge, 2025; Simon et al., 2015) and suggests that, within the pleasantness-sensitive mesolimbic system (Sabatinelli et al., 2007), instructed outcome prediction is a key organizing dimension of neural response. Meta-analytic evidence further supports this interpretation: across 50 Monetary Incentive Delay studies, NAc and insula are reliably recruited during anticipation (Oldham et al., 2018), and an fMRI meta-analysis distinguishing wanting (cue-predicted reward) from needing (homeostatic food value) localizes wanting-related activity specifically to VTA, striatum, and pallidum (Bosulu et al., 2022).

Two alternative accounts warrant consideration. First, a large literature argues that monetary, erotic, food, and drug rewards engage a common mesolimbic network (Sescousse et al., 2013), raising the question of why in this study Food+ cues elicited a stronger NAc response than erotica. However, those conclusions are drawn largely from comparisons across studies, each sampling only a narrow portion of the reward space. Second, a related concern is that mesolimbic responses may rescale to the local stimulus context (Cox & Kable, 2014; Khaw et al., 2017; Padoa-Schioppa, 2009), such that the response to a cue depends partly on the alternatives presented alongside it. Our within-session design speaks directly to both issues: when outcome-predictive and non-predictive food cues are presented together with arousal-matched emotional images, NAc preferentially tracks outcome prediction rather than appetitive visual content. A direct test of the rescaling account would require varying the stimulus set across experiments, an important target for future work.

### Learned outcome associations modulate visual processing

Outcome associations also modulated visual cortex. Food+ elicited greater visual cortex activation than Food−, despite the two conditions consisting of visually identical photographs. This finding converges with our earlier ERP results showing that food cues predicting real delivery elicit larger late positive potentials (LPP) than non-predictive food cues (Versace et al., 2019), and with source analyses implicating occipital and parietal visual cortex as the primary generators of that contingency-driven LPP enhancement (Sabatinelli et al., 2006). The present fMRI results provide convergent spatial evidence that learned outcome associations modulate neural responses in visual cortex. Thus, outcome prediction does not appear simply to add a subcortical motivational signal on top of an otherwise unchanged sensory processing, but also to modulate the sensory response to the cue itself. This interpretation is consistent with evidence that reward history modulates early visual processing. Reward-predictive stimuli capture attention involuntarily (Anderson et al., 2011), and value signals penetrate to spatially selective areas of visual cortex including V1 (Serences, 2008). This modulation appears to be supported by a midbrain–visual cortex pathway (Hickey & Peelen, 2015) and by fronto-parietal coupling with NAc during reward-biased attention (Padmala & Pessoa, 2011). Our Food+ > Food− visual cortex effect extends this reward-driven modulation account by showing that sensory modulation can arise from a learned cue–outcome contingency rather than a rewarded target. One plausible circuit mechanism is top-down feedback from reward and attention networks (Gottfried et al., 2003; Vuilleumier et al., 2001), but the present data cannot directly test that possibility because standard connectivity analyses cannot distinguish feedback-driven modulation from shared task-evoked activity. We therefore treat this as a hypothesis rather than a demonstrated result.

### Temporal dissociation of anticipatory and consummatory signals

Comparing the Food+ Yes > No cue-period contrast with the Juice Yes > No outcome-period contrast revealed a clear anatomical and temporal dissociation consistent with the wanting/liking framework (Berridge & Kringelbach, 2015). This interpretation should be viewed as inferential rather than definitive, however, because the present study did not collect subjective hedonic ratings of the juice. During the cue period, NAc showed the strongest anticipatory signal, differentiating Food+ trials that participants later accepted from those later declined, with no significant outcome-period effect. This pattern identifies NAc as the region showing the clearest anticipatory wanting effect in our ROI set, consistent with its established role in incentive salience attribution (Berridge & Robinson, 2003) and anticipatory reward encoding (Knutson et al., 2001; Oldham et al., 2018). During the outcome period, vmPFC showed the strongest effect, tracking whether juice was delivered. Interpreting this signal as consummatory liking is supported by prior work linking vmPFC activity to subjective hedonic ratings of received rewards (Bartra et al., 2013; Chib et al., 2009; Clithero & Rangel, 2014; Kringelbach, 2005; Plassmann et al., 2010), although that inference rests on that broader literature rather than on direct behavioral measurement in the present sample. The larger outcome-than cue-period effect in vmPFC further suggests that this region is more strongly engaged by realized than anticipated value, consistent with its proposed role as a value accumulator (Kable & Glimcher, 2009). Dorsal anterior insula participated significantly in both phases with comparable effect sizes, consistent with its proposed role as an interoceptive hub integrating motivational salience during anticipation with the sensory-affective qualities of the received outcome (Critchley et al., 2004; Kleckner et al., 2017; Seeley et al., 2007).

This interpretation aligns with translational evidence that wanting and liking are pharmacologically dissociable in humans: naltrexone, an opioid antagonist, selectively reduces self-reported wanting and modulates frontostriatal connectivity, whereas dopamine blockade produces no comparable effect (Soutschek et al., 2021). This dissociation complements rodent evidence for anatomically distinct NAc circuits supporting incentive wanting and hedonic liking (Costa et al., 2016; Smith & Berridge, 2007; Wassum et al., 2011), as well as meta-analytic evidence separating cue-triggered wanting-related activation in striatum, VTA, and pallidum from homeostatic needing-related activation in mid-insula and caudal-ventral putamen (Bosulu et al., 2022). Together this literature supports interpreting the present cue-period NAc effect as an anticipatory signal consistent with wanting.

The Yes > No cue-period NAc signal can also be read as a Pavlovian-to-instrumental transfer (PIT) effect, in which a learned Pavlovian predictor (Food+) invigorates the instrumental response of accepting the juice. Human fMRI studies have localized general PIT, the nonspecific invigoration of reward-seeking by reward-predictive cues, to NAc (Prevost et al., 2012; Talmi et al., 2008), consistent with rodent evidence that phasic NAc dopamine release tracks reward seeking during PIT expression (Wassum et al., 2013). NAc PIT signals also predict relapse in alcohol dependence (Garbusow et al., 2016). Thus, the Food+ cue-period NAc activation observed here is consistent with general PIT circuitry (Cartoni et al., 2016) and suggests that the Yes > No contrast captures the same motivational process that drives cue-triggered approach in addiction models.

### Representational geometry is organized by outcome prediction

RSA provided convergent, multivariate evidence that outcome prediction was the dominant organizing principle of the neural response across the full ten-region network. Of the four models tested, only the Food Reward model significantly predicted the observed representational geometry, with Food+ representationally isolated from all other conditions. Most notably, Food+ and Food− showed the greatest pairwise dissimilarity in the matrix, despite being the only condition pair matched for visual content.

This pattern becomes more understandable when one distinguishes quantitative from qualitative differences across regions. Across much of the network (visual cortex, mPFC, IFG, amygdala, insula subdivisions), differences between emotional and non-emotional conditions were largely quantitative: Erotica and Mutilations evoked larger responses than Neutral and Food, but the overall pattern across regions remained broadly similar (Sabatinelli et al., 2005; Sambuco, Bradley, Herring, & Lang, 2020; Versace et al., 2014). In the mesolimbic reward system, by contrast, Food+ produced a qualitatively different pattern, with NAc responding strongly to Food+ but weakly to the other conditions (Knutson et al., 2001; Sabatinelli et al., 2007; Sambuco, Bradley, Herring, & Lang, 2020). Because RSA captures the shape of profiles across regions rather than absolute response magnitudes (Kahnt, 2018; Kriegeskorte et al., 2008), Food+ emerges as the representational outlier. This approach has shown that representational geometry can reveal organizational principles not evident from univariate activation alone, including identity-specific reward codes in lateral OFC that are functionally dissociable from value-general codes in vmPFC and are dynamically updated by selective satiation (Howard et al., 2015; Howard & Kahnt, 2017), Thus, learned outcome associations did not override emotional representation; rather they reorganized the geometry of the network by recruiting a distinct subcortical circuit that responds selectively to cues predicting real outcomes (Haber & Knutson, 2010).

### Individual differences and future directions

A notable feature of the paradigm is the substantial between-subject variability in juice acceptance. This variability reflects genuine individual differences in appetitive motivation under conditions of real juice delivery, and, importantly, enabled the within-subject wanting versus liking contrasts. The present sample, characterized by elevated mean BMI and substantial variability in acceptance behavior, provides a strong foundation for future individual-difference analyses linking neural wanting signals to real-world eating, consistent with prior MID-fMRI work linking individual differences in striatal reward anticipation to socioeconomic and developmental factors(Mullins et al., 2020). Prior work has shown that striatal responsivity to food cues prospectively predicts weight gain (Demos et al., 2012; Stice et al., 2010; Stice & Yokum, 2016), and a meta-analysis of 45 studies estimates a medium effect of food-cue reactivity on eating and weight outcomes (r ≈ 0.33) (Boswell & Kober, 2016). The obesity literature is nevertheless heterogeneous: a preregistered meta-analysis found no uniform group difference between obese and lean adults in food-cue responses, with age moderating effects in the insula and fusiform gyrus (Morys et al., 2023). This nuance motivates prospective, within-subject designs like the one introduced here, which probe individual-level NAc reactivity under real juice delivery. We (Versace et al., 2016, 2019) identified electrophysiological reactivity profiles that predict subsequent eating behavior, and converging behavioral work in humans has shown that individuals who attribute stronger incentive salience to reward-predictive cues exhibit larger Pavlovian-to-instrumental transfer effects (Degni et al., 2022; Garofalo & Di Pellegrino, 2015). The present fMRI paradigm can be understood as the neuroimaging analogue of that approach: by embedding real gustatory outcomes within an emotionally rich visual context, it is designed to increase sensitivity to neural differences between individuals for whom food-predictive cues hold strong versus weak motivational significance. Future studies with larger samples could test this correspondence directly, combine the present paradigm with LPP-based cluster classification in the same participants, or use pharmacological manipulations (e.g., GLP-1 receptor agonists) to ask whether cue-evoked wanting signals are modified by interventions targeting appetitive motivation.

Exploratory analyses further revealed associations between pre-scan satiety and anterior insula and amygdala responses to food cues. Although these effects did not survive correction for multiple comparisons, they are consistent with the proposed interoceptive function of these regions (Craig, 2009; Critchley et al., 2004; Kleckner et al., 2017) and suggest that future studies with larger samples and repeated satiety assessments could examine whether interoceptive sensitivity modulates food cue processing independently of metabolic hunger state.

### Conclusion

Food cues that predict a real gustatory outcome selectively recruited nucleus accumbens, were anatomically and temporally dissociable from outcome-period signals in vmPFC, and dominated the multivariate representational geometry of motivationally relevant stimuli, even in the presence of highly arousing emotional images. Whether a visual stimulus engages mesolimbic reward circuitry depends more on what it predicts than what it depicts. These findings establish a paradigm for investigating how wanting and liking signals are disrupted in conditions of dysregulated appetitive motivation, and for identifying neural endophenotypes that render individuals differentially vulnerable to cue-triggered behavior.

## Acknowledgments

This work was supported by grant R01DA032581from the National Institute on Drug Abuse (PI: FV), by grant P30CA016672 from the National Cancer Institute to the University of Texas MD Anderson Cancer Center, and by generous philanthropic contributions to the Cancer Neuroscience Program of the University of Texas MD Anderson Cancer Center.

